# Nest defense intensity does not vary with time of day in American Herring Gulls (*Larus smithsonianus*)

**DOI:** 10.1101/2025.06.13.659635

**Authors:** Shailee S. Shah, Maddie E. Ellms, Frances Dygean, Scarlett Gunasekera, Emily Hart, Reka E. Ivanyi, Morgan Lane, David Liu, DeAnna Pitcher, Abigail Saucier, Kathleen F. Schroeder, Joseph Triquet, Kit M. Williams, Kristen M. Covino

**Affiliations:** Cornell Lab of Ornithology, Ithaca, NY, USA; Ecology and Evolutionary Biology Department, Cornell University, Ithaca, NY, USA; College of Life Sciences and Agriculture, University of New Hampshire, Durham, NH, USA; Biology Department, Loyola Marymount University, Los Angeles, CA, USA; College of Arts and Sciences, Cornell University, Ithaca, NY, USA; College of Agriculture and Life Sciences, Cornell University, Ithaca, NY, USA; Biology Department, Environmental Studies Program, Middlebury College, Middlebury, VT, USA; Southern Maine Community College, South Portland, ME, USA; Shoals Marine Laboratory, University of New Hampshire, Durham, NH, USA; Gulls of Appledore Research Group, Appledore Island, ME, USA

**Keywords:** American Herring Gull, antipredator defense, diurnal pattern, *Larus smithsonianus*, simulated predatory threat

## Abstract

Nest defense in birds is vital for the protection of their young but can prove energetically costly. Birds often show plasticity in nest defense depending on factors such as threat level, mate presence, nest stage, and body condition. Such factors can vary over temporal extents ranging from a diel cycle to a breeding season or an individual’s lifetime. Quantifying diurnal variation in nest defense intensity can be particularly useful for understanding time-energy budgets in breeding birds, but it remains understudied. Here we investigated how nest defense by American Herring Gulls (*Larus smithsonianus*) in response to a predatory threat varies with time of day. We simulated predatory threat at 26 Herring Gull nests during four different times of day: “Early Morning” (0530 – 0600 h), “Late Morning” (1000 – 1030 h), “Afternoon” (1400 – 1430 h), and “Evening” (1900 – 1930 h). Based on previous findings of diurnal activity patterns in American Herring Gulls and closely related species, we predicted that nest defense intensity—measured as aggressiveness of response and latency to calm—would be greater during the early morning and late evening than at other times. Contrary to our prediction, we did not detect evidence of difference in nest defense intensity among the four sampled time periods. We found a marginal effect of the presence of the non-incubating mates on the incubating gulls’ likelihood of moving off the nest, but their response scores and latency to calm were not significantly different than when the non-incubating mate was absent. Previous work has shown that nest defense intensity in American Herring Gulls is low overall during the incubation stage which could explain our results.

## Introduction

Nest defense is defined as “a behavior that decreases the probability that a predator will harm the contents of the nest (eggs or chicks) while simultaneously increasing the probability of injury or death to the parent” (Montgomerie and Weatherhead 1988). As such, nest defense is a costly behavior which can vary in intensity depending on various factors that influence its relative costs and benefits (Caro 2005). Previous studies across a variety of avian species have shown that the intensity of nest defense varies based on internal factors such as parental sex (Brunton 1990; Reyer et al. 1998; Galeotti et al. 2000; Paredes et al. 2006; Kryštofková et al. 2011; Tkaczyk et al. 2022) and experience (Knight and Temple 1986; Thornhill 1989) as well as external factors such as offspring age and vulnerability (Regelmann and Curio 1983; Redondo and Carranza 1989; Pavel and Bureš 2001; Brown and Brown 2004; Crisologo and Bonter 2017; Strnadová et al. 2018), presence and behavior of mate (Chase 1980; Crisologo and Bonter 2017), and mating system (Larsen 1991). However, though birds often show distinct diurnal cycles in activity patterns (Robbins 1981), few studies have examined how nest defense varies with time of day (Burger 1981; Bukacińska and Bukaciński 1994).

Birds show distinct diurnal patterns in activities such as territorial behavior (Galusha and Amlaner 1978; Burger 1981; Bukacińska and Bukaciński 1994; Jahn et al. 2017; Moran et al. 2019; Staicer et al. 2019; Searfoss et al. 2020), foraging (Kelly and Wood 1996; Rastogi et al. 2006; Regular et al. 2010; Dismas et al. 2021; Capilla-Lasheras et al. 2024), movement and habitat use (Galusha and Amlaner 1978; Harmange et al. 2021; Klarevas-Irby and Farine 2024), incubation (Drent 1970; Rastogi et al. 2006), mate guarding and copulation (Burger 1976; Galusha and Amlaner 1978), and antipredator responses (Metcalfe and Ure 1997; Ferguson et al. 2019). These diurnal activity patterns are in response to diurnal variation in factors such as energetic condition (Metcalfe and Ure 1997; Ferguson et al. 2019), circulating hormone levels (Ramenofsky et al. 1999; Schwabl et al. 2016; Eikenaar et al. 2020), ambient temperature (Brown and Brown 2004), and activity of predator and prey species (Drent 1970; Galusha and Amlaner 1978; Regular et al. 2010; Harmange et al. 2021; Ribic et al. 2021).

Although not as well studied as in other groups, such factors may be particularly important in colonial nesting seabirds. Seabirds often forage many miles away from their nest, experience diurnal variation in prey availability, and face nest predation pressure from other individuals in the colony (Schreiber and Burger 2002). For example, in a Black Skimmer (*Rynchops niger*) colony, intraspecific aggression levels showed a daily biphasic pattern, peaking in the early morning and late afternoon in conjunction with peaks in bird activity on the colony level (Burger 1981). Similarly, in a Black-headed Gull (*Larus ridibundus*) colony, intraspecific aggression peaked in the late afternoon to coincide with increased colony activity as birds departed to and arrived from foraging sites (Bukacińska and Bukaciński 1994).

Seabirds are also widely studied in their breeding colonies as indicator species for marine ecosystems and are facing severe declines worldwide (Piatt et al. 2007; Phillips et al. 2023). Understanding fine-scale variation in their nest defense patterns is thus vital to understanding shifts in their time-energy budgets (Viera et al. 2011; Ponchon et al. 2014). However, to our knowledge, no study has examined variation in nest defense with time of day using an experimental approach.

Here, we examined diurnal patterns in nest defense intensity in nesting American Herring Gulls (*Larus smithsonianus*). Although now considered a separate species, the closely related European Herring Gull (*Larus argentatus*) is classically known to exhibit a suite of nest defense behaviors including vocalizations, aggressive displays and postures, and engagement in physical altercations with predators that vary with nest defense intensity (Tinbergen 1961). American Herring Gulls also demonstrate plasticity in nest defense behaviors under different cost-benefit scenarios, varying their intensity of defensive behavior in relation to perceived threat level from auditory (MacLean and Bonter 2013; Shah et al. 2015) and visual (Covino et al. 2023) cues, and in response to offspring vulnerability (Crisologo and Bonter 2017). American and European Herring Gulls have also been shown to exhibit diurnal activity patterns in nest attendance (Morris 1987) as well as individual and colony-level rest and activity patterns (Galusha and Amlaner 1978; Hayward et al. 2009), suggesting that factors impacting nest defense intensity may vary with time of day in this species.

Our study addresses the gap in research on diurnal patterns in American Herring Gull nest defense behavior. We experimentally simulated predatory threat at nests of incubating adults at different times of day and quantified nest defense intensity. Previous work on closely related Larid species has shown that activity in colonies peaks in the early morning and late in the day (Galusha and Amlaner 1978) and that agonistic behavior is positively correlated with colony activity (Burger 1976; Conover and Miller 1980; Burger 1981; Bukacińska and Bukaciński 1994). In our study population, the main nest predators are other American Herring Gulls and co-nesting Great Black-backed Gulls (*Larus marinus*) (MacLean and Bonter 2013). Thus, we predicted that nest defense intensity would similarly vary with colony activity, being most intense in the early morning and evening.

## Methods

### Study system

Our study site was Appledore Island, Maine, USA (42.9891° N, 70.6142° W) which is home to a mixed species colony of approximately 500 American Herring Gulls and 300 Great Black-backed Gulls. We conducted our study on 26 American Herring Gull nests, at all of which the parents were already incubating, with complete clutches (*N* two eggs = 2, *N* three eggs = 24). We selected focal nests that were relatively isolated from one another and that did not have a nearest neighbor within 10 m to prevent a confounding response from neighboring birds from interfering with the trial. We conducted trials from 4 – 9 June 2024 when local sunrise and sunset times were approximately 0504 h and 2019 h EDT respectively. The short sampling period purposefully controlled for variation in intensity of nest defense, as well as other daily activity patterns, at different stages in the nesting cycle (Crisologo and Bonter 2017; Włodarczyk 2017; Elderbrock et al. 2025). To investigate diurnal variation in nest defense, we conducted trials during four time periods: “Early Morning” (0530 h – 0600 h), “Late Morning” (1000 h – 1030 h), “Afternoon” (1400 h – 1430 h), and “Evening” (1900 h – 1930 h). We did not conduct trials in the event of rain to avoid interrupting incubation in inclement weather. In the seven cases when the first egg started hatching prior to completing all trials, we ceased conducting trials on that nest since nest defense shows a sharp increase at hatch day (Crisologo and Bonter 2017). Thus, our trial sample sizes for response score varied across the four time periods: *N* = 23 early morning, *N* = 25 late morning, *N* = 21 afternoon, *N* = 21 evening.

### Field methods

Humans have historically posed a threat to nesting gulls by harvesting both eggs and adult birds for food (Drury 1973), and continue to pose a threat by using frightening devices such as pyrotechnics (Belant 1997), and even illegal lethal techniques (Samusenko and Pyshko 2023), to actively discourage nesting gulls. Additionally, American Herring Gulls at the same study site have been shown to perceive a direct walk towards the nest as a greater threat than a tangential walk (Burger and Gochfeld 1981). Thus, for each trial, a person, the “intruder”, simulated a predator by walking toward the focal nest, standing 1 m from the nest for 30 s, and then walking away from the nest. The trial was considered started as soon as at least one parent from the focal nest was able to visually perceive the intruder and considered ended when the intruder was no longer visible to either parent. The intruder always walked at a steady pace with a neutral posture and expression while looking at the ground to minimize variation in perceived threat across trials since human gaze can impact perceived threat level (Goumas et al. 2020). Intruders wore standardized clothing (neutral-colored and long-sleeved) and a black or gray bicycle helmet (for human safety) to avoid adding confounding variables (Covino et al. 2023).

Nest defense intensity was scored using a numerical scale (Table 1; modified from Covino et al. 2023; MacLean & Bonter 2013; Shah et al. 2015) as the most intense response that was exhibited by the initially-incubating gull throughout the 30 s test period. A second person, the “observer”, positioned themselves far enough away from the focal nest where their presence did not disturb the gull and independently scored the nest defense intensity. The trial only commenced when the focal gull showed no signs of vigilance. After 30 s the intruder walked away from the nest and the observer recorded the amount of time taken for the initially incubating gull to return to its previous state before the trial began (*i.e.*, incubating without any defensive behavior). This measure of latency to calm was not recorded if the observer could not clearly see the gull return to the nest and was tracked for a maximum of 300 s (latency to calm sample sizes; *N* = 22 early morning, *N* = 24 late morning, *N* = 20 afternoon, *N* = 19 evening). For each focal nest, the order in which the time-of-day trials were conducted was randomized and each focal nest experienced each of the four time of day trials only once. Additionally, the intruder was also randomized across all trials and 8 total intruders, such that no nest experienced the same intruder twice.

**Table 1.** Ethogram to rate the behavioral response of American Herring Gulls. The response score increases with intensity of aggression and energy expenditure, modified from (Covino et al. 2023).

| Response Score | Response Description |
| --- | --- |
| 0 | No response |
| 1 | Initial increase in vigilance in response to the intruder's presence, followed by relaxation |
| 2 | Sustained vigilance for entire trial; neck slightly raised, looking at intruder, but does not interrupt activities (*e.g.*, panting, preening) |
| 3 | Sustained vigilance for entire trial; neck fully outstretched, looking at intruder, activities often interrupted, use of '*kek-kek*' calls either while still incubating or while moving away from the intruder |
| 4 | Extreme vigilance for at least a portion of the trial, and overall vigilance sustained for entire trial; use of '*yeow*' calls in combination with vigilance (above-described behaviors) and/or oblique posture while still incubating or while moving away from the intruder |
| 5 | Extreme vigilance for at least a portion of the trial, and overall vigilance sustained for entire trial; use of above-described behaviors in combination with interruption of incubation, standing up off eggs (over the nest), and/or oblique posture, or with interruption of prior positioning |
| 6 | Extreme vigilance sustained for entire trial; use of above-described behaviors in combination with movement no closer than 0.5 m of the intruder including localized charging or lunging towards (typically without contact) intruder and/or sustained oblique posture |
| 7 | Extreme vigilance sustained for entire trial; use of above-described behaviors in combination with movement closer than 0.5 m of the intruder, including charging towards intruder, attempts at some physical contact (*e.g.* biting at legs), and/or short hopping flights towards the intruder |
| 8 | Extreme vigilance sustained for entire trial; use of above-described behaviors in combination with direct and intense attack behaviors, including swooping flight, sustained attempts of physical contact |

Though we also recorded if the non-incubating mate was present within observable distance from the nest, we did not use its response score in analyses. Thus, in trials (*N* = 26) where both mates were present, the response score representing that trial was that of the initially incubating gull. In all cases when the non-incubating mate was also present, it always participated in joint nest defense with the incubating mate. Instances where the incubating gull left the nest and the area completely, without any other response, were excluded from analyses (5 out of 90 [5.6%] trials). Only two of these trials were from the same nest, there was no consistent pattern of time of day, and the non-incubating mate was not present for four of the five trials when this occurred. Additionally, though we did not include these five trials in our main analyses, upon exploration, we also found that including them in the analyses with a response score of 0 did not change the results (see *SI*).

### Statistical analysis

To investigate the effect of time of day on nest defense intensity we fit two mixed-effects models with response score (ordinal error structure) and latency to calm (gaussian error structure) as dependent variables and time of day as a categorical fixed effect. We also included mate presence (yes/no) as a fixed effect since the presence of the non-incubating mate has been shown to affect nest defense intensity in this study population (Crisologo and Bonter 2017). The ordinal model had a logit link function, and the proportional odds assumption was tested with a Brant-Wald test (see *SI*). Latency was cube root transformed for normality. We added date and nest ID as random effects since we conducted multiple trials at each nest and multiple trials per day. However, since date did not explain any variance, we removed it from the final model.

To further investigate if the incubating bird changes its nest defense behavior in response to the presence of its mate, we ran a *post-hoc* analysis to examine if the presence of their mate changed the likelihood that incubating gulls would get off their nest during the predation simulation trial. We fit a generalized linear mixed-effects model with a binomial error structure, with whether the incubating mate got off the nest (yes = 1, no = 0) as the dependent variable and mate presence (yes/no) as the categorical fixed effect. Date and nest ID were included as random effects. All analyses were performed in R version 4.2.2 (R Core Team 2022). Models were fit using the R packages “ordinal” (Christensen 2026) or “lme4” (Bates et al. 2015). Where applicable, we used the “Anova” function to run a type II ANOVA for all models. Variance inflation factor, which indicates multicollinearity between fixed effects (Fox and Monette 1992), was < 2 (no multicollinearity) for models with multiple fixed effects.

## Results

Response scores of incubating American Herring Gulls to a simulated predatory threat ranged from 2 – 8 (*i.e.*, sustained vigilance for entire trial to swooping and direct aerial attacks towards the intruder) with a mean (± SD) of 5.47 (± 1.87) (Fig. S1A). Latency to calm for the incubating parent ranged from 0 – 300 s (with 300 s being our cutoff), with a mean (± SD) of 48.00 (±79.76) s (Fig. S1B). Latency to calm only exceeded 300 s for < 6% (5 out of 85) of the trials. Response score and latency to calm were moderately correlated (Spearman’s ρ = 0.39; Fig. S2).

We found that American Herring Gulls did not exhibit variation in nest defense intensity with time of day in terms of both response score (χ²_3,85_ = 3.32, *P* = 0.34) or latency to calm (*F*_3, 40.91_ = 0.57, *P* = 0.64) (Fig. 1, Tables S1, S2). Presence of the non-incubating mate was also not associated with nest defense intensity (response score: χ²_1,85_ = 1.17, *P* = 0.28; latency to calm: *F*_1, 61.08_ = 0.04, *P* = 0.85; Tables S1, S2), though it had a marginally significant positive association with the incubating gull’s likelihood of moving off the nest during the trial (Z = 1.78, *P* = 0.08, 95% CI = -0.18 – 3.64). However, since the presence of the non-incubating mate covaried significantly with nest ID (chi-square test: χ²_24,85_ = 38.60, *P* = 0.03) such that some nests always or never had the non-incubating mate present (Fig. S3), the effect of mate presence could be partially confounded by the random effect of nest ID and should thus be interpreted with caution.

**Figure 1.**
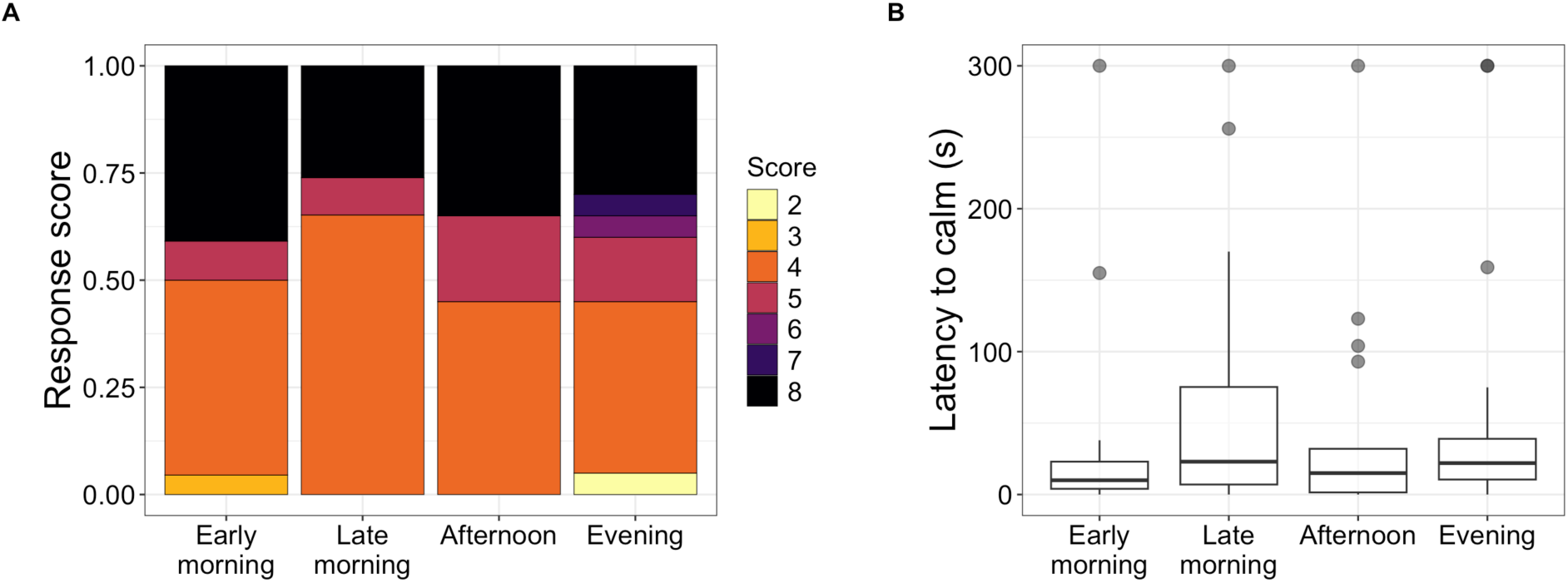
American Herring Gulls incubating at nests did not exhibit variation in (A) Response score (*N* = 26 nests, 85 trials; colors represent ordinal response scores) or (B) Latency to calm (*N* = 25 nests, 80 trials) across four different times of day (x-axis). Response score was measured based on an ethogram of response aggressiveness and intensity to a potential predator in American Herring Gulls (Table 1), and latency to calm was measured as time in seconds taken for the focal bird to return to its pre-trial state (*i.e.*, incubating without any defensive behavior), with a maximum cutoff at 300 s. Both behavioral measures were recorded in response to a simulated predatory threat of a human walking up to and standing for 30 s next to an American Herring Gull incubating a nest. **Alt text.** Multipaneled figure. Panel A on the left shows the proportion of American Herring Gulls exhibiting each response score (range: 2 – 8) to a simulated predatory threat across four categorical times of day as a stacked bar chart. Panel B on the right shows the latency to calm following the same trials across four categorical times of day as boxplots.

## Discussion

Since nest defense can be energetically costly, and even prove lethal, birds often adjust the intensity of nest defense behavior based on both intrinsic (*e.g.*, body condition) and extrinsic (*e.g.,* threat level) factors (Montgomerie and Weatherhead 1988). Such factors may vary with time of day (Drent 1970; Galusha and Amlaner 1978; Conover and Miller 1980; Robbins 1981; Kelly and Wood 1996), thus impacting the intensity of nest defense behavior. Here, we investigated whether nest defense in American Herring Gulls, a colonial ground-nesting seabird with a suite of nest defense behaviors that have been shown to vary with perceived threat (MacLean and Bonter 2013; Shah et al. 2015; Crisologo and Bonter 2017; Covino et al. 2023), varies with time of day. We found no evidence that nest defense intensity varies with time of day in our study population on Appledore Island, Maine. Additionally, incubating gulls did not exhibit a more intense defense response when the non-incubating mate also participated in nest defense. We detected a trend of incubating mates changing their nest defense behavior when the non-incubating mate was present, but this result should be interpreted with caution. Overall, our results suggest that nest defense in American Herring Gulls does not vary with time of day, and though previous studies have found an effect of joint nest defense, the association may be more nuanced than previously considered.

Nest defense has been shown to vary with time of day in some avian species including seabirds such as Black Skimmers (Burger 1981) and Black-headed Gulls (Bukacińska and Bukaciński 1994), though not in a consistent manner. Whereas nest defense in Black Skimmers peaked early in the morning (Burger 1981), Black-headed Gulls showed a peak in the late afternoon, the latter likely associated with increased movement of birds to and from foraging sites (Bukacińska and Bukaciński 1994). Contrary to both results, we found no variation in nest defense intensity with time of day in American Herring Gulls. However, whereas previous studies were based on observations of conspecific and heterospecific interactions, we used an experimental approach with human presence as a simulated predatory threat. The nesting stage could also have influenced our result, since previous work in the same population found that nest defense intensity of American Herring Gulls is lowest overall at the incubation stage (Crisologo and Bonter 2017). Additionally, our study largely included nests in areas frequented by humans where American Herring Gulls nest in a looser colony structure (Savoca et al. 2011) to ensure that experimental trials would not be subject to interference from birds at neighboring nests. Habituation to close human presence could thus explain the lack of variation in nest defense intensity with time of day in response to a simulated predatory threat, though previous work in our study system found no evidence of habituation (MacLean and Bonter 2013; D. Titmuss unpubl. data).

We found that nest defense intensity did not vary with the presence of the non-incubating mate who always participated in joint nest defense. Though a previous study in the same population found that overall nest defense intensity was higher when both parents were present, they considered the total response of all gulls participating in nest defense (Crisologo and Bonter 2017). Thus, our results suggest that individual responses of American Herring Gulls to predators may not be flexible enough to compensate for the absence of the non-incubating mate, at least during the incubation stage. Additionally, we found that mate presence was not evenly distributed across our trial nests—for some nests the non-incubating mate was present during all four trials whereas for others the non-incubating mate was never present. Though it was beyond the scope of our study to determine the sex or individual identity of the incubating and non-incubating mates, previous work has shown that nests with more equitable distribution of both incubation and presence near nest when not incubating have higher reproductive success in American Herring Gulls and closely related species (Burger 1987; Kavelaars et al. 2019).

We found a marginally significant trend of the incubating gull being more likely to move off the nest during nest defense when its non-incubating mate was participating in joint nest defense. Though, as described above, mate presence was not evenly distributed among the nests, this observation raises questions about coordinated nest defense that can be examined in future studies. For example, both parents may engage in more aggressive nest defense jointly than when defending the nest alone if the presence of their mate ameliorates the risk of nest failure due to injury or death to one parent (Montgomerie and Weatherhead 1988) or as a response to their mate’s “cooperative” behavior (Chase 1980). A more nuanced study of nest defense behavior of each parent singly versus both parents jointly is needed to understand coordination of nest defense behavior in species with biparental care.

Overall, our results suggest that multiple factors can simultaneously influence nest defense intensity. Though we did not find an association of nest defense intensity with time of day or presence of the non-incubating mate at the clutch stage, future studies should examine how these associations vary at different nest stages and with variation in coordination of offspring care duties between the parents in this and other species with biparental care.

## Acknowledgements

We thank the Shoals Marine Laboratory for giving us access to Appledore Island and its facilities and the staff and faculty for their support. We thank Dylan Titmuss for their help with our experimental design, Mary Elizabeth Everett and Sarah J. Courchesne of the Gulls of Appledore Research Group for project support, and Mitchell Norris for data collection in the field.

## Permits and ethics protocols

Ethical approval for this research study was granted by the Shoals Marine Lab and University of New Hampshire IACUC (Approval number 210503).

## Author contributions

Conceptualization: all authors; Data curation: all authors; Formal analysis: S.S.S. and K.M.C.; Funding acquisition: all authors; Investigation: all authors; Methodology: all authors; Project administration: S.S.S. and K.M.C.; Resources: S.S.S. and K.M.C.; Software: S.S.S.; Supervision: S.S.S. and K.M.C.; Validation: all authors; Visualization: S.S.S.; Writing – original draft: all authors; Writing – reviewing & editing: S.S.S. and K.M.C. contributed to project development, at least one component of the study itself, data management, and draft composition. SSS and KMC supervised the project development, data collection, analyses, manuscript preparation, and edited and revised the final version of the manuscript.

## Generative AI

The authors did not use Generative AI in the production of this article.

### Data and code availability

Data and code are available on GitHub at https://github.com/shailee93/diurnal_effects_joint_nest_defense_herring_gulls

