## Supplementary information for "Nest defense intensity does not vary with time of day in American Herring Gulls (*Larus smithsonianus*)"

This file includes:
Supplementary Text
Figures S1-S3
Tables S1-S2

**Supplementary Text**

*Results including trials where the incubating gull left the nest and area completely*

Retaining the five trials where the incubating gull left the nest and area completely, without any other response, did not change our results. These trials were assigned a response score of 0. Nest defense intensity did not vary significantly with time of day or presence of their non-incubating mate in terms of both response score (time of day: $\chi^2_{3,90} = 3.36$, $P = 0.32$; mate presence: $\chi^2_{1,90} = 1.09$, $P = 0.30$) or latency to calm (time of day: $F_{3,\ 57.79} = 1.51$, P = 0.22; mate presence: $F_{1,\ 69.02} = 0.03$, $P = 0.86$).

*Checking model assumptions for the ordinal model*

We used the function "brant" from the R package *brant* (Schlegel and Steenbergen 2020) to perform a Brant-Wald test to test the assumption of proportional odds. Since only one trial each had a response score of 2 and 3, those outcome categories were too sparse for a reliable test statistic. We therefore excluded these two observations solely for the purpose of conducting the Brant-Wald test. The omnibus Brant–Wald test did not indicate a violation of the proportional-odds assumption, $\chi^2_{12} = 9.44$, $P = 0.66$. Similarly, none of the predictor-specific tests were statistically significant (all $P > 0.05$).

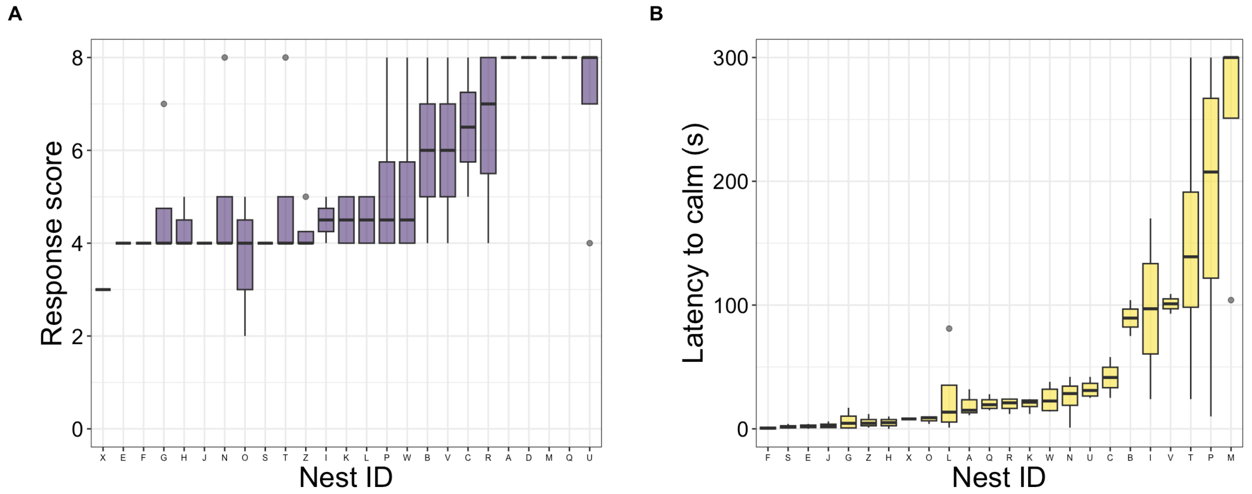


**Figure S1.** Variation in (A) Response score (*N* = 26 nests) (0: least to 8: most aggressive and energy intense response, see Table 1 for details), and (B) Latency to calm (*N* = 25 nests), or time taken for the focal bird to return to its pre-trial state (*i.e.,* incubating without any defensive behavior) of American Herring Gulls at nests to a simulated predatory threat, with a maximum cutoff at 300 s. Both behavioral measures were recorded in response to a simulated predatory threat of a human walking up to and standing for 30 s next to an American Herring Gull incubating a nest. Each nest (x-axis) experienced the simulated predator threat up to four times total to test for an effect of time of day ("Early Morning": 0530 – 0600 h, "Late Morning": 1000 – 1030 h, "Afternoon": 1400 – 1430 h, "Evening": 1900 – 1930 h) across six days. Nests are arranged in order of median response score or latency to calm (lowest to highest) for visual purposes.

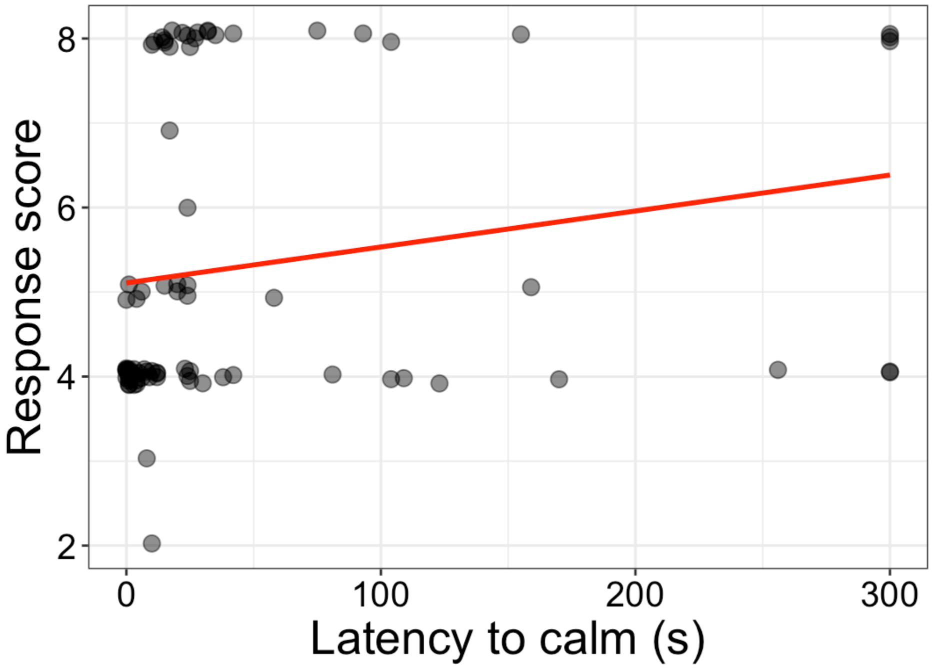


**Figure S2.** The two metrics of nest defense intensity, response score and latency to calm, were moderately correlated across trials (*N* = 80). Points indicate raw data slightly jittered vertically to improve visual discrimination.

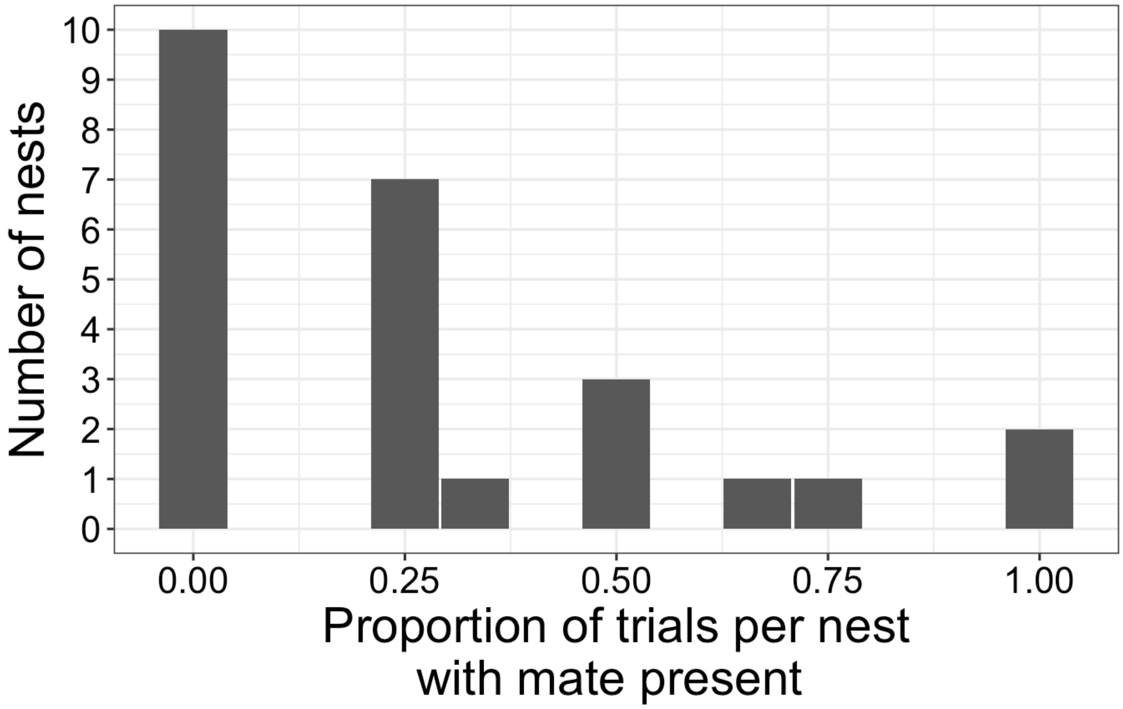


**Figure S3.** Presence of the non-incubating mate was not randomly distributed among the trials conducted at our focal nests. Two nests, for example, consistently had the mate present for all four trials (*i.e.*, proportion of trials per nest with mate present = 1.00), whereas more than a third of the nests ($N$ = 10), never had the mate present in the vicinity for any of the trials (*i.e.*, proportion of trials per nest with mate present = 0.00).

**Table S1.** Results of mixed-effects model with an ordinal error structure estimating the effects of time of day (TOD; "Early Morning" (0530 h – 0600 h), "Late Morning" (1000 h – 1030 h), "Afternoon" (1400 h – 1430 h), and "Evening" (1900 h – 1930 h)) and presence of non-incubating mate (yes/no) on response score of incubating American Herring Gulls during trials of simulated predatory threat ($N$ = 26 nests, 85 trials). Random effects of nest ID was included.

| **Fixed effect** | **Estimate** | **Std. Error** | ***Z*** | ***P*** | **95% CI** |
|---|---|---|---|---|---|
| TOD: Late Morning | -1.17 | 0.73 | -1.60 | 0.11 | -2.60, 0.26 |
| TOD: Afternoon | -0.03 | 0.71 | -0.04 | 0.97 | -1.42, 1.36 |
| TOD: Evening | -0.32 | 0.72 | -0.45 | 0.65 | -1.73, 1.09 |
| Mate Presence: Yes | 0.72 | 0.67 | 1.08 | 0.28 | -0.59, 2.04 |

**Table S2.** Results of linear mixed-effects model estimating the effects of time of day (TOD; "Early Morning" (0530 h – 0600 h), "Late Morning" (1000 h – 1030 h), "Afternoon" (1400 h – 1430 h), and "Evening" (1900 h – 1930 h)) and presence of non-incubating mate (yes/no) on latency to calm of incubating American Herring Gulls during trials of simulated predatory threat (*N* = 25 nests, 80 trials). Random effects of nest ID was included. Latency to calm was cube root transformed for normality and had an upper bound of 300 s, which was only reached in < 6% of trials.

| **Fixed effect** | **Estimate** | **Std. Error** | **df** | ***t*** | ***P*** | **95% CI** |
|---|---|---|---|---|---|---|
| TOD: Late Morning | 0.31 | 0.30 | 54.71 | 1.02 | 0.31 | -0.28, 0.90 |
| TOD: Afternoon | 0.09 | 0.31 | 54.05 | 0.30 | 0.76 | -0.50, 0.72 |
| TOD: Evening | 0.39 | 0.32 | 54.63 | 1.24 | 0.22 | -0.22, 1.01 |
| Mate Presence: Yes | -0.06 | 0.31 | 62.50 | -0.20 | 0.84 | -0.66, 0.53 |
